# Geographic range dynamics drove ancient hybridization in a lineage of angiosperms

**DOI:** 10.1101/129189

**Authors:** R.A. Folk, C.J. Visger, P.S. Soltis, D.E. Soltis, R.P. Guralnick

## Abstract

Factors explaining global distribution patterns have been central to biology since the 19th century, yet failure to combine dispersal-based biogeography with shifts in habitat suitability remains a present-day setback in understanding geographic distributions present and past, and time-extended trajectories of lineages. The lack of methods in a suitable integrative framework stands as a conspicuous shortcoming for reconstructing these dynamics. Here we showcase novel methods to overcome these methodological gaps, broadening the prospects for phyloclimatic modeling. We focus on a clade in the angiosperm genus *Heuchera* endemic to southern California that experienced ancient introgression from circumboreally distributed species of *Mitella,* testing hypotheses regarding biotic contact in the past between ancestral species lacking a fossil record. We obtain strong support for a past contact zone in northwestern North America, resolving this paradox of hybridization between ancestors of taxa currently separated by ∼1300 km.

---

Geographic ranges of montane organisms have been and continue to be dramatically driven by climate change,^1-5^ sometimes resulting in surprising highly disjunct distributions. A remarkable example in plants is the ancestral occurrence of hybridization between taxa that are currently allopatric; past co-occurrence has been hypothesized to be driven by Pleistocene climate change increasing suitable habitat.^6-8^ For extant lineages, these scenarios can be tested using paleoclimatic projections and existing ecological niche modeling (ENM) methods. Yet gene flow has also been posited between species ancestral to clades of extant organisms^9^ – hypotheses for which methodological developments have failed to keep pace. In these instances, the object of inference is the degree of overlap in niche envelope of two lineages that are no longer extant, such that ENM methods must be combined with ancestral niche reconstruction (“phyloclimatic modeling”^10^). If these methods are extended to estimate *distributions* of input variables on ancestral lineages, they can be used to place constraints on the extent of suitable habitat for taxa, or in other words, the ancestral niche envelope. The extent of habitat suitability in ancestral taxa can be assessed directly in multivariate ecological niche space (E-space), for which the only hard limit in terms of temporal depth is the ability to obtain confident ancestral E-space reconstructions under available models. In addition, for time periods for which paleoclimatic data are available, the inferred extents of ancestral habitat suitability can be translated into geographic space (G-space; i.e., “geographic projection”). Development of these key extensions of phyloclimatic modeling—particularly the direct use of nodal variance estimates and geographic projection of nodes—provide an opportunity to test directly not only hypotheses about niche shifts, but also environmental and geographic *overlap* through time, reconstructing the potential for historical biotic interactions.

The flowering plant genus *Heuchera* (Saxifragaceae) has experienced an extreme level of gene flow among lineages, both contemporary and ancestral, making it an excellent test case^9,11-13^. An earlier study^9^ used coalescent simulations that supported a hypothesis of ancient introgression rather than lineage sorting to explain well-resolved but extremely discordant phylogenomic data. The most dramatic of these instances, represented by a disjunction of ∼1300 km, involves the well-supported capture of the chloroplast genome of the ancestor of two species of *Mitella* (extant distribution circumboreal and eastern U.S.) by the ancestor of a clade of five southern Californian species of *Heuchera* (hereafter referred to as “California *Heuchera*”) (Fig. 1a).^9,13^ Species of these two genera are remarkably distinct morphologically (Fig. 1a), yet genera of Saxifragaceae are known to hybridize and can be crossed artificially^9^.

We have developed a computational pipeline to implement novel methods that use distributions of niche space for ancestral lineages and predict their geographic ranges using paleoclimate data. We asked whether the ancestors of *Mitella* and California *Heuchera* (1) overlapped in niche space, and (2) whether and where they overlapped in geographic space under paleoclimate scenarios. We then integrated these estimates with a time-calibrated phylogeny and biogeographic reconstructions to refine further areas of plausible co-occurrence that may have facilitated hybridization. The approach provided here represents a significant advance over previous approaches for general hypotheses about historical range dynamics, providing a toolkit to broaden and generalize the prospects for phyloclimatic modeling.

**Fig. 1.**
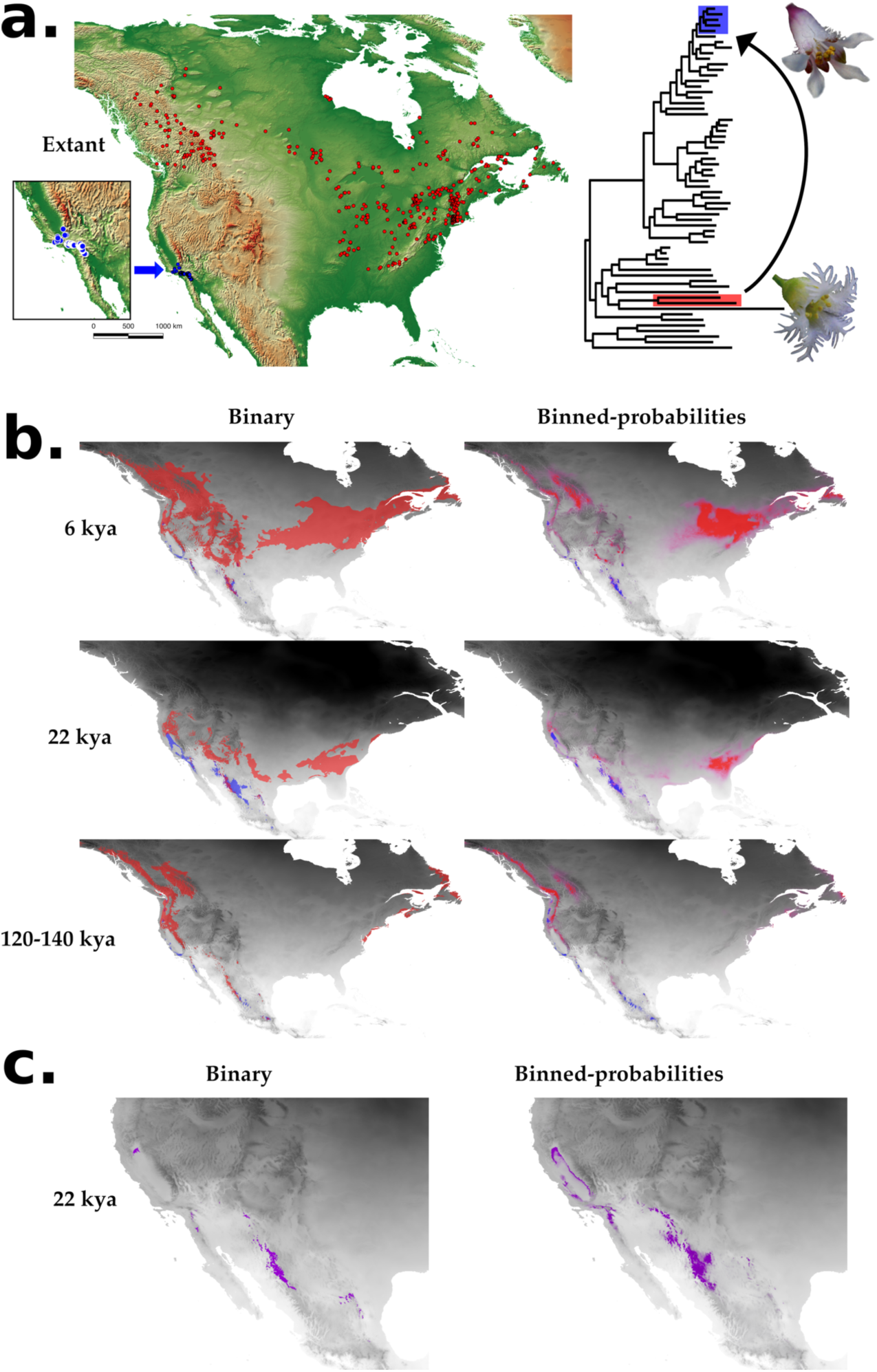
(a.) Extant distribution of *Mitella* (red) and California *Heuchera* (blue), and a phylogeny plotting the focal clades and inferred introgression event. (b.) Ancestral geographic projections of estimated niche space for ancestors of *Mitella* (red) and California *Heuchera* (blue), at all three time slices with both projection methods (c) Predicted LGM (22 kya) intersection of ranges between *Mitella* and *California* with both projection methods in purple; zoomed to predicted overlap in western North America. These projections show narrow-buffer models; wide-buffer model projections are given in Supplemental Fig. S6. Last interglacial = 120-140 kya, LGM = 22 kya, Holocene Optimum = 6 kya.

**Fig. 2.**
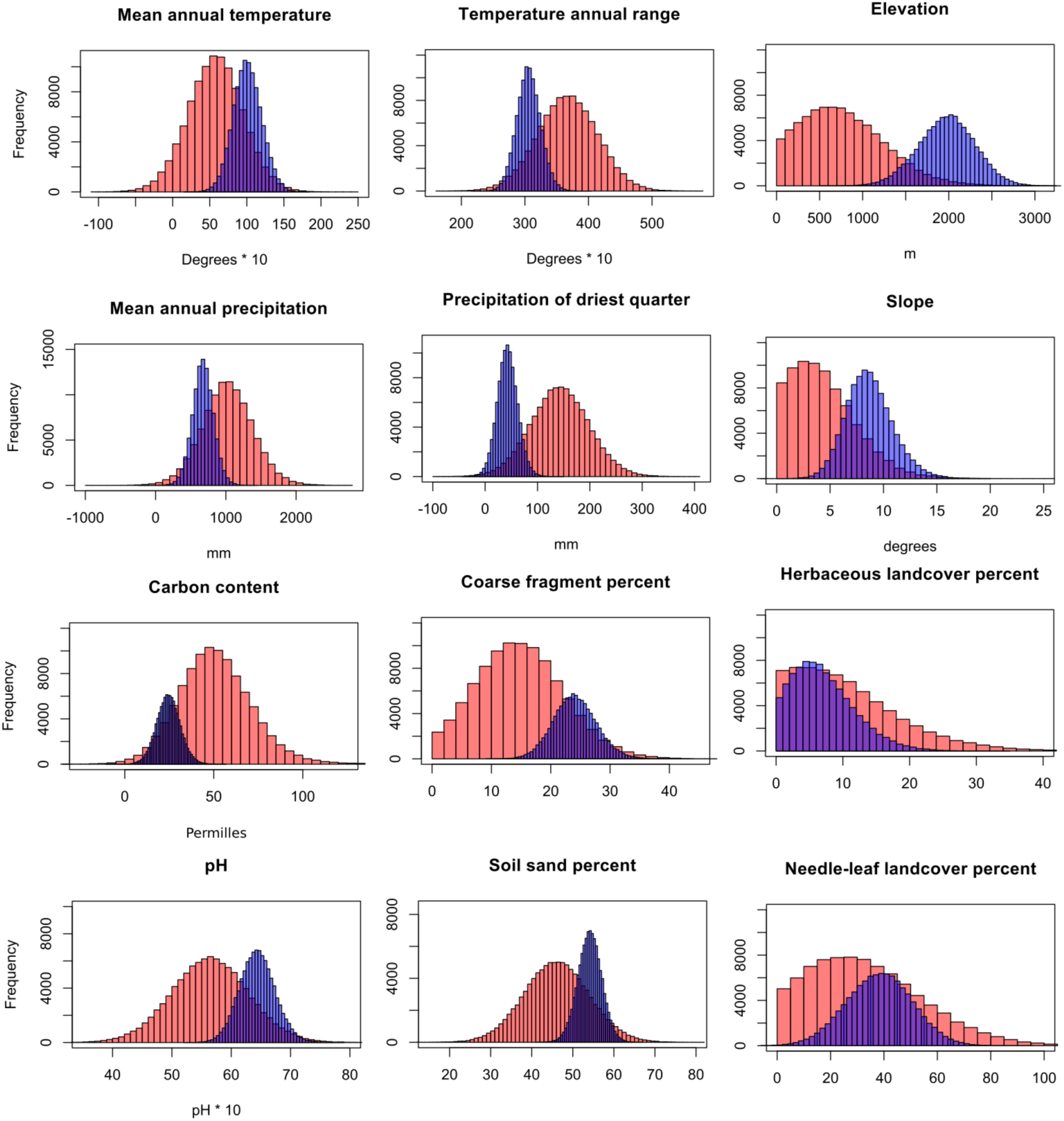
Histograms summarizing pooled MCMC distributions across all 12 variables examined for ancestors of *Mitella* (red) and California *Heuchera* (blue). The y-axis represents absolute sample number. Narrow-buffer models were used for these histograms.

## RESULTS

### Ancestral niche reconstruction

Ancestral reconstructions of environmental parameters (Figs. 2 and 3) showed phylogenetic structure (Fig. 3). As expected under a Brownian model (cf. ^14^), relative error (nodal icons, Fig. 3) was approximately normal and highest at the earliest nodes and internal nodes with long parent and daughter branches; shallow divergences had relatively low uncertainty.

**Fig. 3.**
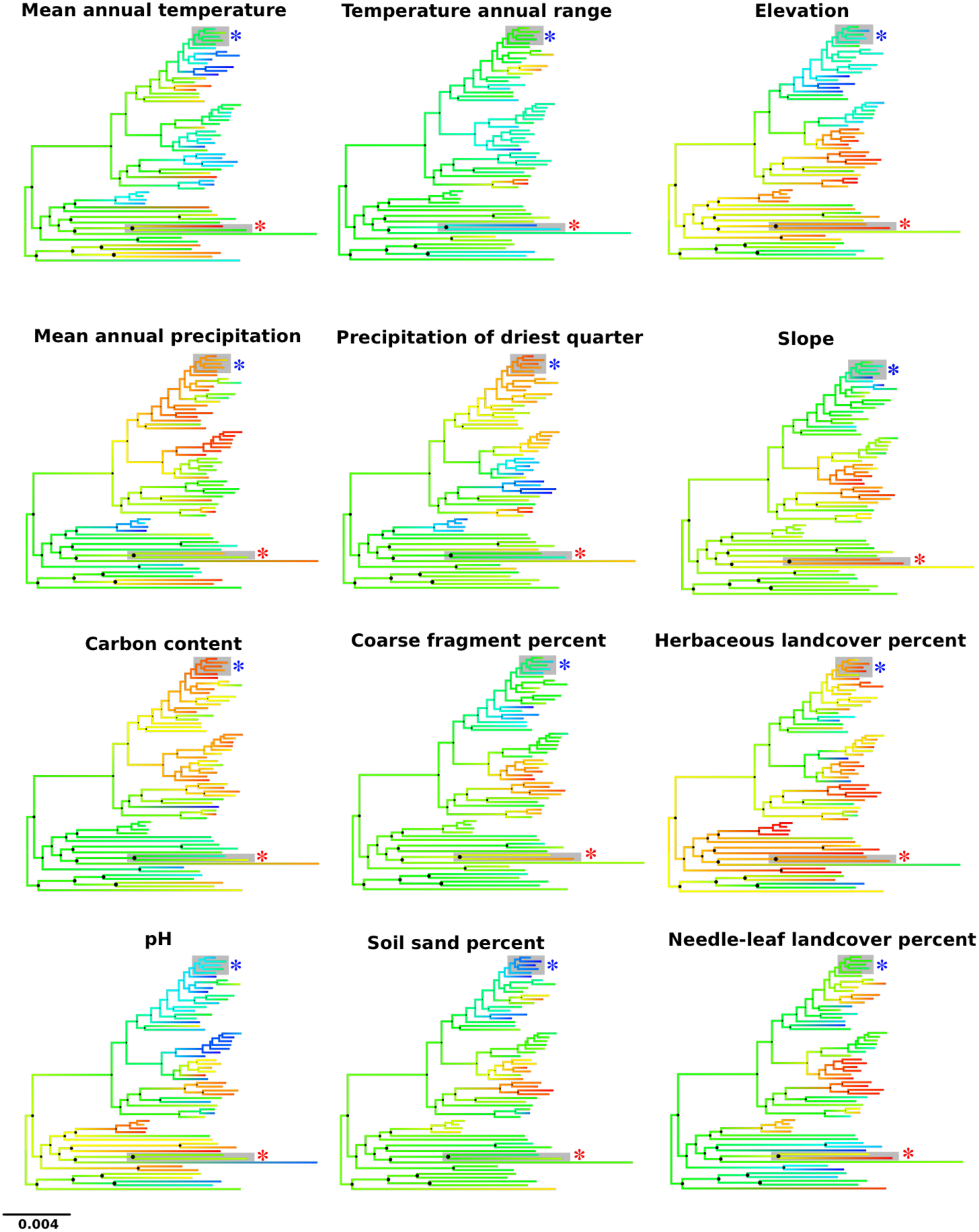
Median values of niche space for all ancestral nodes and tip taxa, using narrow-buffer models. Branches are colored by median values scaled from the minimum observed (red) to the maximum (blue). Black circles at nodes represent by their diameters the relative credibility interval breadth of the nodes. Species labels are not shown for compact representation, but tips follow the exact sequence of Supplementary Fig. S2. Asterisks denote focal clades – blue for California *Heuchera* and red for *Mitella.*

**Table 1.** Quantification of variable-wise environmental (E-space) overlap as intersections of pixel-wise joint posterior probabilities of occurrence between the two taxa (defined in Supplemental Methods).

|  | NARROW-BUFFER<br>MODELS | WIDE-BUFFER<br>MODELS |
| --- | --- | --- |
| Mean annual temperature | 0.4013 | 0.4149 |
| Temperature annual range | 0.3031 | 0.3133 |
| Mean annual precipitation | 0.4083 | 0.3988 |
| Precipitation of driest quarter | 0.1594 | 0.1592 |
| Carbon content | 0.2648 | 0.3252 |
| Coarse fragment percent | 0.3450 | 0.3378 |
| Elevation | 0.1378 | 0.1505 |
| Herbaceous land cover percent | 0.6608 | 0.6594 |
| Needle-leaf land cover percent | 0.6033 | 0.5867 |
| pH | 0.3537 | 0.3539 |
| Soil sand percent | 0.3654 | 0.3677 |
| Slope | 0.3057 | 0.3130 |

**Table 2.** Quantification of geographic (G-space) overlap between the ancestors of *Mitella* and California *Heuchera* as intersections of pixel-wise joint posterior probabilities of occurrence between the two taxa (defined in Supplemental Methods). Under both training region formulations, overlap probability is greatest during glacial conditions (boldface).

|  | Global<br>cumulative overlap | North American<br>cumulative overlap |
| --- | --- | --- |
| Holocene optimum (6 kya) | 0.058290 | 0.005526 |
| Last Glacial Maximum (22 kya) | **0.066441** | **0.008329** |
| Last interglacial (120-140 kya) | 0.019045 | 0.002915 |

WIDE-BUFFER MODELS
|  | Global<br>cumulative overlap | North American<br>cumulative overlap |
| --- | --- | --- |
| Holocene optimum (6 kya) | 0.075261 | 0.007003 |
| Last Glacial Maximum (22 kya) | **0.082419** | **0.010098** |
| Last interglacial (120-140 kya) | 0.025021 | 0.003659 |

### Ancestral overlap

The ecological niche spaces (E-space) of the most recent common ancestors (MRCAs) of *Mitella* and California *Heuchera* have distinct but overlapping distributions for all variables examined (Table 1; Fig. 2). When the ancestral distributions are projected into environmental conditions estimated for the Holocene, LGM, and interglacial periods (Fig. 1b), the geographic distributions of probabilities of suitability are distinct and parapatric. Areas of greatest potential geographic overlap include northern California and the Sierra Madre of Mexico. The cumulative overlap (quantified by the intersection of the joint probability density in G-space, see methods; Table 2) was highest under glacial maximum conditions. This was equally true for models under two training region buffers (next-best timeslice ∼2/3 the predicted range overlap) although the absolute magnitude differed. Overall geographic ranges predicted for both model sets were very similar, but wide-buffered models resulted in somewhat larger ranges.

### Biogeographic inference

AIC and LRT favored DEC+J (Supplementary Figure S4), yet for the focal ancestral taxa, DEC and DEC+J did not differ qualitatively in geographic range inference; this was also largely true of the tree as a whole. The MRCAs of both *Mitella* and California *Heuchera* contain a Pacific region, but have essentially no probability of presence in the Sierra Madre. Hence, when dispersal and vicariance are considered, suitable habitat for these ancestral taxa in the Sierra Madre is ruled out, leaving northern California as the remaining region of highest overlap probability. As previously observed under a parsimony reconstruction,^15^ nodes ancestral to *Heuchera* and other genera contain the Pacific region without exception. The use of the DEC model relaxes the assumption of single-state geographic ranges, yet a substantial portion of nodes ancestral to, and early-diverging within, *Heuchera* are still solely Pacific.

## DISCUSSION

Using our newly developed methods, we demonstrate that cool conditions during the Pleistocene promoted a parapatric distribution that is not presently possible, but was conducive at the time to facilitate gene flow between California *Heuchera* and species of *Mitella.* For all environmental variables examined, the ancestors of California *Heuchera* and *Mitella* had broadly overlapping inferred historical distributions despite being separated by ∼6 my of divergence. Hence, shared habitat suitability during the Pleistocene reasonably supports the existence of opportunities for ancient gene flow between the ancestors of these currently highly disjunct modern taxa. We also asked how ancestral suitability would be geographically distributed under various climate scenarios.

Under a Pleistocene cooling scenario, ancestral suitable range projections were very different from present-day distributions. The suitable range of the MRCA of *Mitella* was much more southerly and narrow than that of the extant species, consistent with typical responses to glaciation for lowland taxa in eastern North America.^16^ The range suitability of the ancestor of the California *Heuchera* clade was much broader than the present range but continuous with it – an expected feature for high-montane organisms in more suitable cooler climates that have sky-island distributions in the present.^17^ Hence, although suitable habitat for the California *Heuchera* clade expanded during the Pleistocene, the pronounced southward shift of *Mitella* at this time compared to the present primarily drove geographic overlap between these clades. Overall, while the suitability prediction for California *Heuchera* oscillated considerably in different time slices, *Mitella* habitat suitability showed more extreme latitudinal shifts, as expected for lowland taxa. Large amounts of suitable climate were inferred for both lineages in the Sierra Madre of Mexico and northern California; suitability in other regions had almost no overlap, suggesting a parapatric pattern of historical contact.

Suitable habitat in Mexico was also detected in present-day models for extant taxa in the California *Heuchera* group. Failure to disperse may explain the current absence of these species in Mexico rather than loss of ancestral suitability. The BioGeoBEARS analysis confirmed this scenario – presence in the Pacific region is supported in the MRCAs of *Mitella s.s*. and California *Heuchera,* and this is true of the sequence of nodes in the phylogeny between these lineages and the root of the *Heuchera* group of genera. The model, however, assigned essentially no probability to presence in Mexico in the MRCA of either clade. This biogeographic result represents the accessible area within which dispersal between predicted suitable habitats is reasonable, suggesting that northern California and not Mexico is the more likely area where ancient gene flow occurred.

Under both cool and warm historical scenarios, we inferred geographic overlap of these ancestral taxa. However, overlap probabilities were much higher during the LGM scenario, particularly compared to interglacial conditions, indicating that cool conditions during the Pleistocene (LGM or prior) would have most greatly facilitated parapatric overlap and produced the greatest opportunity for gene flow.

### New prospects in the projection of ancestral habitat suitability

The methods presented here for ancestral projection are novel in using distributions of ancestral estimates for the purpose of projecting suitability probabilities of MRCAs in geographic space. The approach provided here represents a significant advance over previous statistically problematic approaches (cf. Supplementary Methods), and our study will therefore broaden and generalize the prospects for “phyloclimatic modeling”. The methods are also highly flexible in being compatible with practically any ENM method due to the PNO-sampling approach.^18^ Additionally, while incorporation of the present-day distribution of taxa with respect to environmental variables has been performed before with various approaches,^19,20^ using a Bayesian framework to simultaneously incorporate phylogenetic uncertainty, reconstruction error, and present-day distribution of niche space with respect to individual environmental parameters is novel. A Bayesian framework is ideally suited for research on ancestral trait distribution, estimation of which is unavoidable in questions involving niche space.

Recently the issue of homology has been raised with respect to ancestral niche reconstruction:^18^ are comparisons of environmental parameters between taxa under a trait model logical? Ancestral reconstruction methods treat draws from niche distributions as true trait values; hence, this is a valid question. This is a general challenge in comparative methods and applies to other “traits” that are actually complex wholes of underlying unitary characters (e.g., suites of simple features such as pollination syndromes and reproductive behaviors, multiple loci controlling a trait, and even amino acid translations of DNA data^21^). The former have been termed “composite” characters and contrasted with their unitary components, termed “reductive” characters, which are the traditional object of systematic inference.^22^ There is a long tradition of analyzing such composite characters on trees, necessitated by the fact that very few phenotypic characters are truly reductive; empirically obtained composite character data can be viewed as imperfect but justifiable heuristic estimates of underlying indivisible traits that have in many cases led to useful empirical conclusions. It would be more desirable to measure and draw conclusions about underlying physiology of (the fundamental) niche, perhaps under a model that can also incorporate trait interactions. Yet these physiological traits are largely unknown as well as laborious to attain directly for large numbers of taxa, and are only indirectly and precariously obtained by attempting to deduce fundamental niche from ENMs that may include environmental covariates of factors that represent realized niche.^23^ Even successfully obtaining underlying physiological limits leaves layers of abstraction between the analysis and reductive phenotypic factors, which will typically be multifactorial genetic traits, likely with epigenetic components. We aver that ENMs of extant taxa using standard methods will contain environmental factors co-distributed with these underlying phenotypic traits, often perhaps quite closely; hence, we are likely capturing much of that signal, as much as is possible given globally available environmental data, with a considerable economy of effort that promises to scale well with questions requiring ambitious taxon sampling.

### New prospects for hybridization detection and event contextualization

Our new methods have broad utility in the study of hybridization, a common process in eukaryotic life. The detection of hybridization has largely been an exercise in molecular data analysis,^9,24-29^ with few exceptions^8^. This assertion is *strictly* true of studies assessing ancestral hybridization (that is, gene flow among species that are not extant). Our study suggests that assessing estimates of environmental envelopes is a promising alternative and complementary *validation* procedure. If candidate lineages involved in interspecific gene flow can be identified using molecular data, then estimating niche envelope overlap for these lineages can assess the feasibility of gene flow *a posteriori*. Beyond simple hypothesis validation, these analyses, in combination with time-calibrated trees, can facilitate the identification of geographic regions and climatic conditions in which particular events could have taken place. For most hybrids that have been identified, these parameters are unknown, hindering understanding of typical pathways of formation and factors that enable long-term survival of recombined lineages as compared to their unhybridized relatives.

### Future directions for ancestral habitat reconstruction

The approaches provided here have broad future implications for the inference of ancestral niche reconstruction. While it has been possible to link niche modeling and phylogenetics^10^, development of streamlined implementations of the methods and suitable downstream applications have not followed. Researchers have been limited both by lack of a firm theoretical foundation and a lack of programs enabling post-processing and analysis (beyond mere visualization) of ancestral reconstructions. Future work focusing on explicit integration of biogeography with habitat suitability will facilitate new forms of hypothesis testing relevant to shallow- and deep-scale diversification, particularly in relation to historical climate change. Assessment of ancestral dynamics in geographic space relies on the availability of historical climate layers that to date are available primarily for relatively shallow time scales; however, analyzing shifts in niche space occupancy possesses no hard temporal limits. We hope that the present case study demonstrates the potential power of applications generating distributions of ancestral suitability as compared to the point estimates or ranges under uncertain model formulations that have solely been inferred up to this point, and the integrative role these data have in evolutionary questions. In particular, inference of ancestral distributions enables the direct examination of geographic range dynamics and lineage overlap, offering glimpses of historical-biotic interactions, which are perhaps more elusive than all other factors that biologists have invoked for explaining present-day distributions and organismal properties.

## METHODS

### Phylogenetics

The phylogenetic framework built on previous results^9,30^ by supplementing outgroup sampling with key taxa to strengthen our identification of introgressed lineages and inferences of ancestral states for the focal groups. Briefly, the earlier study sequenced 277 nuclear loci via the targeted enrichment method presented in^30^ (total target length ∼400,000 bp). We sampled 42 of 43 known species of *Heuchera*. To improve outgroup representation, we mined transcriptome resources from the 1KP project (onekp.com/) and generated new genomic targeted sequencing and transcriptome data for 13 additional outgroup taxa, summarized in Supplemental Table S1. Sequencing and alignment followed previously reported methods^9,30^ with modifications noted in Supplemental Methods.

### Species tree estimate

Given the known similarity of results from concatenation and coalescence methods for this dataset^9^, we used the former (methods in ^9^; modifications noted in Supplemental Methods). To summarize state reconstructions on a single optimal tree, we inferred a total-evidence phylogeny in RAxML (using all individuals sampled, and took the best ML tree), pruned to a single individual per species at random. 1000 boostrap replicates on the pruned individual set were used to integrate phylogenetic uncertainty in ancestral estimation. Given our use of a Bayesian reconstruction method, we also made a trial of concatenated Bayesian phylogenetic inference in MrBayes (^31^; not shown); this resulted in optimal or near-optimal (∼1) posterior probability on all nodes for a tree highly similar to our best ML tree, a result that defeats the purpose of incorporating phylogenetic uncertainty and is perhaps consistent with notions that posterior probabilities represent overestimates.^32^

### Chloroplast gene tree estimate

To verify the chloroplast capture scenario^9^ with additional outgroup sampling, we constructed and analyzed an augmented chloroplast matrix with RAxML rapid bootstrapping and best tree search (option -f a) with 5,000 replicates and a GTR-GAMMA model, with coding and noncoding nucleotide positions set as distinct partitions, following methods in ^9^.

### Phylogenetic dating

We verified the compatibility of an approximate Pleistocene timeframe for the lineages of interest with present knowledge by performing secondary dating in BEAST following ^33^, using two prior formulations as noted in Supplemental Methods. Both chloroplast and nuclear data were used to independently estimate the implied timeframe of gene flow.

### Point record synthesis

We synthesized available point records from all taxa in the species tree in major online data repositories (see Supplemental Methods); after assessing sampling weaknesses in point records from these repositories, strategic taxa of *Heuchera* were identified, imaged, and georeferenced from specimen loans. Additional records that were not georeferenced were obtained where necessary from the online repositories noted above, as well as the primary literature, field observations, and herbarium material from our previous work (see Supplemental Methods). We primarily used GEOLocate supplemented by verification in Google Earth and Google Maps, following best practices including, briefly: using only unique and unambiguous place names, discarding localities that refer to large metropolitan areas or cultivated records, verifying that cardinal directions and distances were correctly calculated, and verifying by satellite imagery that broadly suitable habitat was present. Final occurrence counts, deduplicated pixel-wise (30-arcsecond resolution), are given in Supplemental Table S2.

### Ecological niche modeling

We assembled 35 layers at 30-second resolution capturing aspects of climate, soil, land cover, and topography. These comprised 19 well-known temperature and precipitation variables (BIOCLIM; http://www.worldclim.org/bioclim gives the complete list of variables and units), three topographical layers (elevation, GTOPO30; https://lta.cr.usgs.gov/GTOPO30; aspect and slope calculated from this layer in qGIS), seven soil layers (bulk density, coarse fragment percent, clay percent, silt percent, sand percent, pH, organic carbon content, SoilGrids1km; averaged in qGIS across layers at 2.5 cm, 10 cm, 22.5 cm, and 45 cm core depth; https://soilgrids.org/), and six land cover classes (percentage cover of: needle-leaf, evergreen broadleaf, deciduous broadleaf, mixed trees, shrubs, herbaceous; http://www.earthenv.org/landcover). Forthwith, “environmental variables” refer to our climatic and non-climatic factors jointly. We assessed correlation to reduce these to 12 representative layers (cf. Supplemental Methods).

To set up training regions for our models, we combined ecoregional and dispersal distance concepts. We used The Nature Conservancy terrestrial ecoregion dataset (http://maps.tnc.org/gis_data.html; based on^34^), taking those that contained points for the species of interest. To incorporate dispersal, we calculated a convex hull containing all points; in the absence of specific dispersal data from *Heuchera* or its relatives (which are expected to be poorly dispersed; cf. ^12^), we conservatively used a distance of 0.5° for a buffer distance around this hull (60 pixels at 30-second resolution, or ∼42.7 km at 40° latitude). The final training region was taken as the intersection of the kept ecoregions and buffered hull. All calculations were performed in qGIS (http://www.qgis.org/) and calculated per species. Effectively, the training region comprised all biomes from which a species is known, trimmed to 0.5° from the most peripheral occurrences; this is summarized in Supplementary Figure S1. To assess the sensitivity of the results to these “tight” training region definitions, we applied a further 1° buffer to these regions; models and all downstream analyses were performed in duplicate with these two sets of training regions, hereafter referred to as “narrow-buffer” and “wide-buffer” models. Training regions were clipped from global rasters using ENMGadgets (https://github.com/vijaybarve/ENMGadgets).

Maxent was run with a maximum of 5,000 iterations (which was generally not exceeded, given an imposed convergence threshold of 0.00001), extrapolation of extant species’ geographic projections (only used for QC) was disabled, and points with some missing data were allowed (the soil layers had missing data due to the way in which they were calculated, although only a few study species had any occurrences with missing values). For all modeled taxa, 25% of the data were set aside for model testing and the remainder were used for training. Models were averaged over 10 bootstrap replicates. To evaluate model performance, we calculated jackknifes and response curves as well as the default ROC and omission plots; model statistics are given in Supplemental Table S2. Models were inferred for 30-second-resolution data; a trial of 2.5-minute models (not shown) resulted in poorer fit statistics.

### Predicted niche occupancy profiles and ancestral reconstruction

To calculate the distribution of niche space occupancy in the models, we calculated predicted niche occupancy (PNO) profiles (following ^35^, for all retained layers, using R package phyloclim^36^). A PNO profile is a density histogram that represents an integration of a single environmental layer (in this case, but not necessarily, one of those used to construct the model) over the suitability surface predicted by an ecological niche model. The result represents the discrete estimate of a probability density function where the environmental variable is the predictor and the function describes probability of suitability. For taxa with fewer than 12 training points, PNO profiles were generated directly from empirical data following methods given in Supplemental Methods.

To incorporate this statistical distribution of extant species niche space occupancy into ancestral reconstruction, we sampled the environmental value of each of these bins proportionally (similar to ^19,35^) for 1,000 samples. Ancestral reconstruction was performed in BayesTraits 2.0 (http://www.evolution.rdg.ac.uk/BayesTraits.html), an MCMC implementation of ancestral reconstruction that allows for directly estimating the distribution of ancestral values. Each sample was taken as the true point value for each extant taxon in a single ancestral reconstruction; this process was repeated for each input variable, resulting in 12,000 ancestral reconstructions. To incorporate phylogenetic uncertainty, ancestral reconstruction was run on a sample of 1,000 RAxML fast bootstraps (option -f a) from a sequence matrix that included one randomly selected individual per species; results were plotted on the optimal phylogenetic estimate (best ML tree with all individuals, pruned to the same randomly selected samples). For both the model and the reconstruction step, MCMC chains were run for 5,000,000 generations, preceded by 5,000,000 generations of burn-in. All priors were uniform over [-300, 5000], the approximate range of environmental data. For tree summary statistics, the median, 2.5^th^, and 97.5^th^ percentiles were calculated for every node in each MCMC chain; these were then averaged across MCMCs to summarize average median and credibility intervals across MCMCs. Output was formatted in BEAST-style extended NEXUS format for visualization. Because more variable intronic data were excluded, not all clades were highly supported; we directly incorporated phylogenetic uncertainty by searching over the 1,000 bootstrap trees with branch lengths in the MCMC analysis.

### Ancestral suitability overlap

We examined ancestral suitability overlap for the two ancestral taxa of the *Mitella s.s*. and California *Heuchera* clades in both E-space and G-space. For E-space, we quantified the intersection^37^ of probability densities (using probability density histograms in 50 equal bins from the minimum to maximum of observed values) of the two MRCAs for each variable to evaluate the relative impact of the environmental factors we used.

**Fig. 4.**
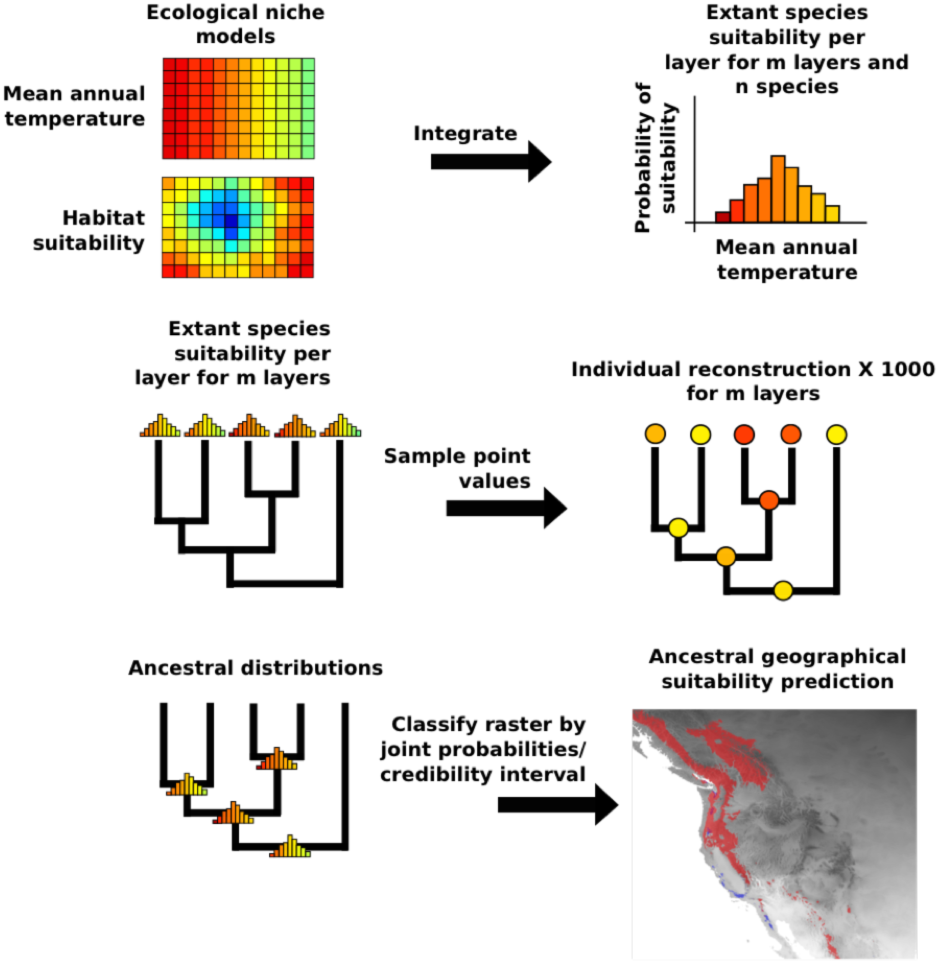
Conceptual summary of the estimation of ancestral niche space performed in 

~~~
ambitus.
~~~ To assess where this range overlap might have been distributed geographically in the past, we considered climatic layers from the Last Glacial Maximum (LGM), mid-Holocene, and last interglacial (available at http://www.worldclim.org/paleo-climate1). For the former two we used the CCSM4 model at 2.5-minute resolution and for the latter, that from ^38^, sampled to 2.5- minute resolution with GDAL, averaging pixels. Historical data are generally not available at both high spatial and temporal resolution, nor is phylogenetic dating to this level of resolution often possible. Hence, LGM data were used as representative of a general Pleistocene cool period; the other two layers were used as two alternative historical-warm scenarios. Of variables used to model niche space, all four retained BIOCLIM layers were available for these periods. We projected nodal variable distributions into geographical space using (1) a binary approach based on an n-dimensional hypervolume comprising credibility intervals, and (2) a binned-probabilities approach that directly estimates pixel-wise joint posterior probabilities. Geographic overlap predictions under the three paleoclimate scenarios were quantified by intersections of joint posterior probability densities; raster method details are given in Supplemental Methods.

### Pipeline construction

We automated the steps outlined in the previous two sections (summarized in Fig. 4) by designing a data analysis pipeline in Python, called 

~~~
ambitus,
~~~

 that serves as a wrapper for BayesTraits, to handle the thousands of ancestral reconstructions required to incorporate PNOs from extant taxa, parallelizing the computationally intensive reconstructions, and automating summary statistics and tree plotting. Some initial data formatting is required (e.g., variable and taxon label standardization) but otherwise operation is straightforward. The program (available at https://github.com/ryanafolk/ambitus/) is compatible with Linux and OS-X and allows customization of a large number of analysis parameters as well as optionally producing extensive diagnostic MCMC output. We also provide shell scripts to perform ancestral geographic projection and overlap quantification.

### Biogeography

Estimates of ancestral biogeographic states were performed in the R package BioGeoBEARS^39^ under both the DEC and the DEC+J model, using our dated nuclear tree under the truncated normal prior. Biogeographic regions were coded broadly from ecoregion concepts, but conceived more broadly to include the main montane physiographic regions that correlate with *Heuchera* distributions in a manageable number of states for the DEC model. Six states were recognized: Pacific-montane (coastal ranges, Pacific NW, Sierra Nevada, Transverse Ranges); Sierra Madre; Basin and Range; Rockies; Eastern U.S.; and Temperate East Asia (this coding scheme is congruent with a previous biogeographic analysis^15^; Supplemental Figure S5). Two further major regions present in the data, Great Plains and Himalayas, were classified as the nearest neighbors of Eastern U.S. and Temperate East Asia, respectively, because only one taxon occurred in each. The maximum allowed ancestral range size was 6 (among extant taxa, the largest was 4). The two models were compared by AIC and likelihood ratio tests (LRT).

### Data availability

All raw sequence data have been deposited in SRA; all DNA assemblies, phylogenetic trees, occurrences, Maxent suitability layers, and ancestral reconstructions have been deposited in Dryad.

### Code availability

Scripts to reproduce ancestral reconstruction analyses are available at GitHub (https://github.com/ryanafolk/ambitus).

## ACKNOWLEDGMENTS

The authors thank H. Owens, N. Barve, and J. Allen for discussions on ENM methods, concepts, and layer sources, B. Drew and M. Gitzendanner for discussions on missing data, C.A. McCormick (NCU) and the curators of the herbaria noted in Supplementary Methods for assisting with occurrence record acquisition, B. Stucky for discussions on Brownian motion and statistical sampling, J.V. Freudenstein for advice concerning homology, E.B. Sessa for reading the paper, the 1KP project for granting access to several transcriptomes, Y. Okuyama for sharing DNA materials of Japanese *Mitella,* the Florida Museum of Natural History for supporting R.A.F. as a visiting researcher, and the NSF PRFB and DDIG programs (DBI 1523667, DEB 1406721) for funding.

## AUTHOR CONTRIBUTIONS

R.A.F., R.P.G., P.S.S., and D.E.S. designed the study; R.A.F. and C.J.V. performed sequencing and occurrence curation; R.A.F. implemented methods and performed analyses, R.A.F., C.J.V., P.S.S., D.E.S., and R.P.G. contributed substantially to the manuscript.

## MATERIALS & CORRESPONDENCE

Correspondence and materials requests should be directed to R.A. Folk.

